# Optimal TELSAM-Target Protein Linker Character is Target Protein-Dependent

**DOI:** 10.1101/2025.08.29.672704

**Authors:** Maria Jose Pedroza Romo, Alihikaua Keliiliki, Jacob C. Averett, Joseph F. Gonzalez, Ethan Noakes, Elijah W. Wilson, Conrad Smith, Blake Averett, Dalton Hansen, Riley Nickles, Miles Bradford, Sara Soleimani, Tobin Smith, Supeshala Nawarathnage, Prasadika Samarwickrama, Ariel Kelsch, Derick Bunn, Cameron Stewart, Wisdom Abiodun, Evan Tsubaki, Seth Brown, Tzanko I. Doukov, James D. Moody

## Abstract

Fusing a variant of the sterile alpha motif domain of the human translocation ETS leukaemia protein (TELSAM) to a protein of interest has been shown to significantly enhance crystallization propensity. TELSAM is a pH-dependent, polymer-forming protein crystallization chaperone which, when covalently fused to a protein of interest, forms a stable, well-ordered crystal lattice. However, despite its success, a challenge persists in that crystal quality and diffraction limits appear to be heavily dependent on the choice of linker between TELSAM and the protein of interest, with identification of a functional linker relying on trial-and-error methods. Likewise, previous studies revealed that the 10xHis tag at the TELSAM N-terminus can either facilitate or hinder the ordered crystallization of target proteins attached via flexible or semi-flexible linkers. To address these challenges, we designed multiple constructs with several types of linkers—rigid (helical fusion), semi-flexible (Pro-Ala_n_), and flexible (poly-Gly)—of varying lengths to fuse a designed ankyrin repeat protein (DARPin) to the TELSAM C-terminus. Semi-flexible and flexible linker constructs were made with and without the 10xHis tag. Our findings indicate that short semi-flexible and rigid linkers consistently yield large crystals within 24 hours with a DARPin target protein, but that flexible linkers perform best with a TNK1 UBA domain target protein. Removing the 10xHis tag enhanced crystallization rates, improved crystal morphology, and increased the crystallization propensity of semi-flexible and flexible linker constructs. While removing the His tag did not have a significant effect on crystal size, it improved the diffraction limits and crystal quality of the 1TEL-PA-DARPin construct. These results suggest that the ideal linker selection primarily depends on the properties of the target protein. Our data support the recommendation to use a short yet flexible or semi-flexible linker between TELSAM and the target protein to facilitate protein crystallization and high-resolution structure determination.

**Synopsis:** In this study, we examine the effect of short to medium-length flexible, semi-flexible, and rigid linkers on the crystallization of a DARPin fused to the 1TEL protein crystallization chaperone, demonstrating that while rigid linkers impair crystallization and reduce diffraction quality, the ideal linker character remain target-protein dependent.

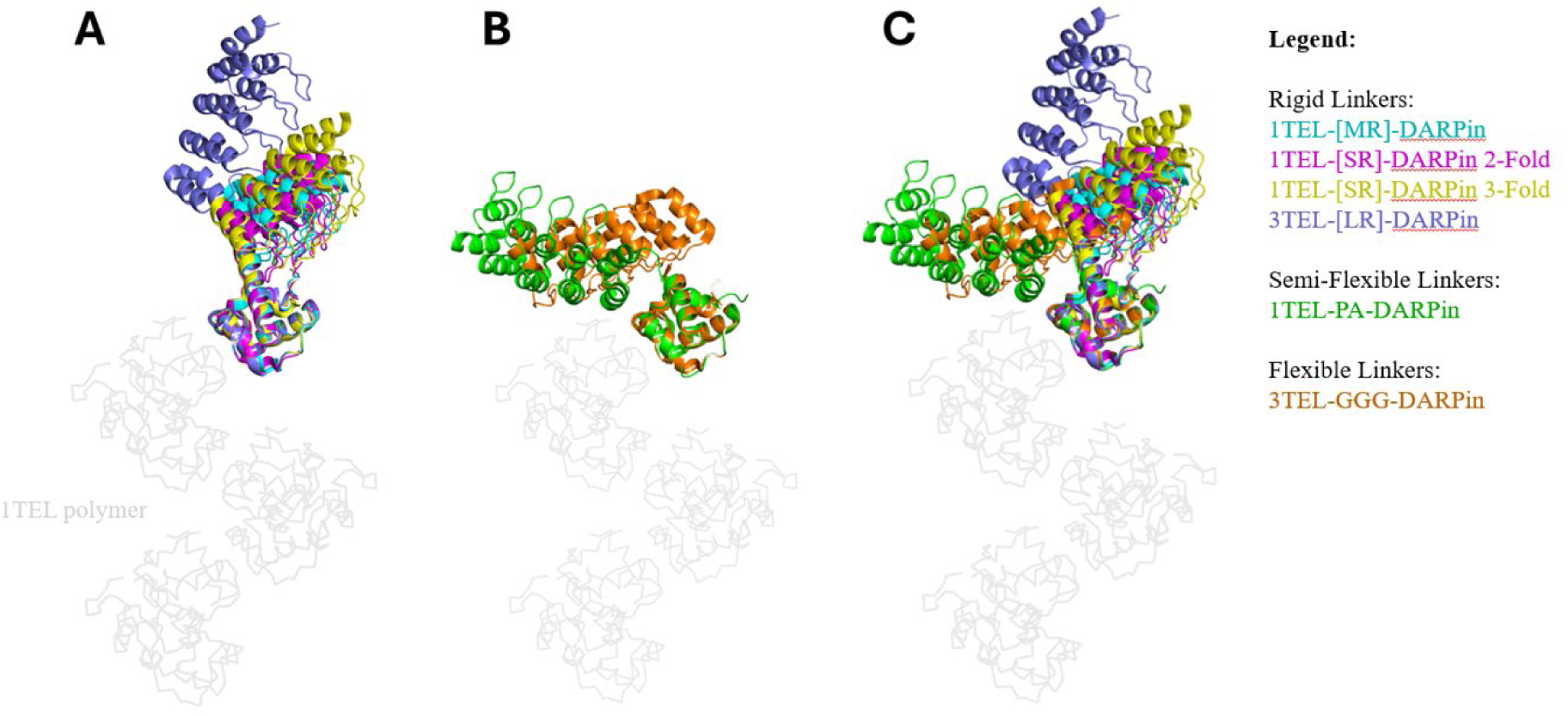

## 1. Introduction

Atomic-resolution macromolecule structures are vital for experimental structural characterization, which is essential for drug and target discovery and design, drug fragment virtual drug screening, and elucidation of protein structure-function relationships in health and disease (Ferreira et al., 2015; Lionta et al., 2014; Wei and McCammon, 2024) (Maveyraud and Mourey, 2020). X-ray crystallography remains the method of choice for obtaining high-resolution structures of macromolecules smaller than 50-100 kDa. Despite advancements in molecular cloning, protein production and purification, crystallization screening, synchrotron data collection, and structure determination software, obtaining diffraction-quality crystals remains the rate-limiting step in characterizing macromolecular structures through X-ray crystallography (Cooper et al., 2011; Dale et al., 2003; Terwilliger et al.)

To address this bottleneck, researchers have turned to innovative approaches, including the use of engineered fusion proteins to enhance crystallization efficiency (Derewenda, 2010). One of these is the human Translocation ETS Leukaemia (TEL, ETV6) sterile alpha motif domain (TELSAM). A point mutation of the TELSAM motif causes it to be soluble at pH 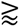 8 and to polymerize into well-ordered helical polymers at pH 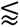 8. These polymers exhibit six TELSAM subunits per complete turn of the helix. Target proteins can be genetically fused to TELSAM, which promotes crystallization by ordering many copies of the target protein along ordered TELSAM polymers, which associate to form the crystal. TELSAM Fusion Crystallization (TFC) offers many advantages in addressing classical protein crystallization challenges: TFC (1) decreases the entropic cost of crystal nucleation and growth; (2) increases crystallization propensity, reproducibility, and speed; (3) enables crystallization at unusually low protein concentrations; (4) yields crystals that reach the same diffraction limits as crystals from traditional methods, but with significantly fewer crystal contacts; and (5) does not perturb target protein structure. To date, TFC has been observed to achieve a 90% success rate in forming crystals (Nauli et al., 2007), promising a highly cost-effective approach for accelerating structural determination. (Gajjar et al., 2023; Nawarathnage et al., 2022; Nawarathnage et al., 2023)

We previously created four protein constructs that fused this same DARPin (McFedries et al., 2013; Seeger et al., 2013) to 1TEL or 3TEL using either a long α-helical fusion or a Gly-Gly-Gly linker (Nawarathnage et al., 2022). 1TEL refers to the use of one TELSAM unit per covalently fused target-protein while 3TEL refers to three TELSAM units covalently fused in tandem with one copy of the target protein fused to the C-terminus of the third TELSAM unit. All constructs gave crystals, but only crystals of the 3TEL constructs were of diffraction-quality. We questioned why the 1TEL constructs failed to yield large or diffracting crystals, but did not pursue the question further at the time. In a later study (Gajjar et al., 2023) we discovered the importance of short rigid linkers in 1TEL-Capillary Morphogenesis Gene 2 (CMG2) von Willebrand Factor Type A (vWA) constructs. Constructs with bulkier, less flexible linkers consistently yielded higher-resolution diffraction data, leading us to hypothesize that optimizing the linkers in 1TEL-DARPin constructs could enable them to form larger, well-ordered crystals. Additionally, another study (Kottur et al., 2022) demonstrated that Pro-Ala and Pro-Ala-Ala linkers provided high-resolution structures of the SARS CoV2 nsp14 methyltransferase domain when using TFC. Herein, we report our work optimizing the 1TEL-DARPin linkers, testing three types of linkers with varying lengths and residue identities: flexible linkers (GG and GGG), semi-flexible (PA and PAA), and rigid α-helical fusion ([SR], [MR], and [LR]) linkers, as in **Table 1**.

**Table 1.**
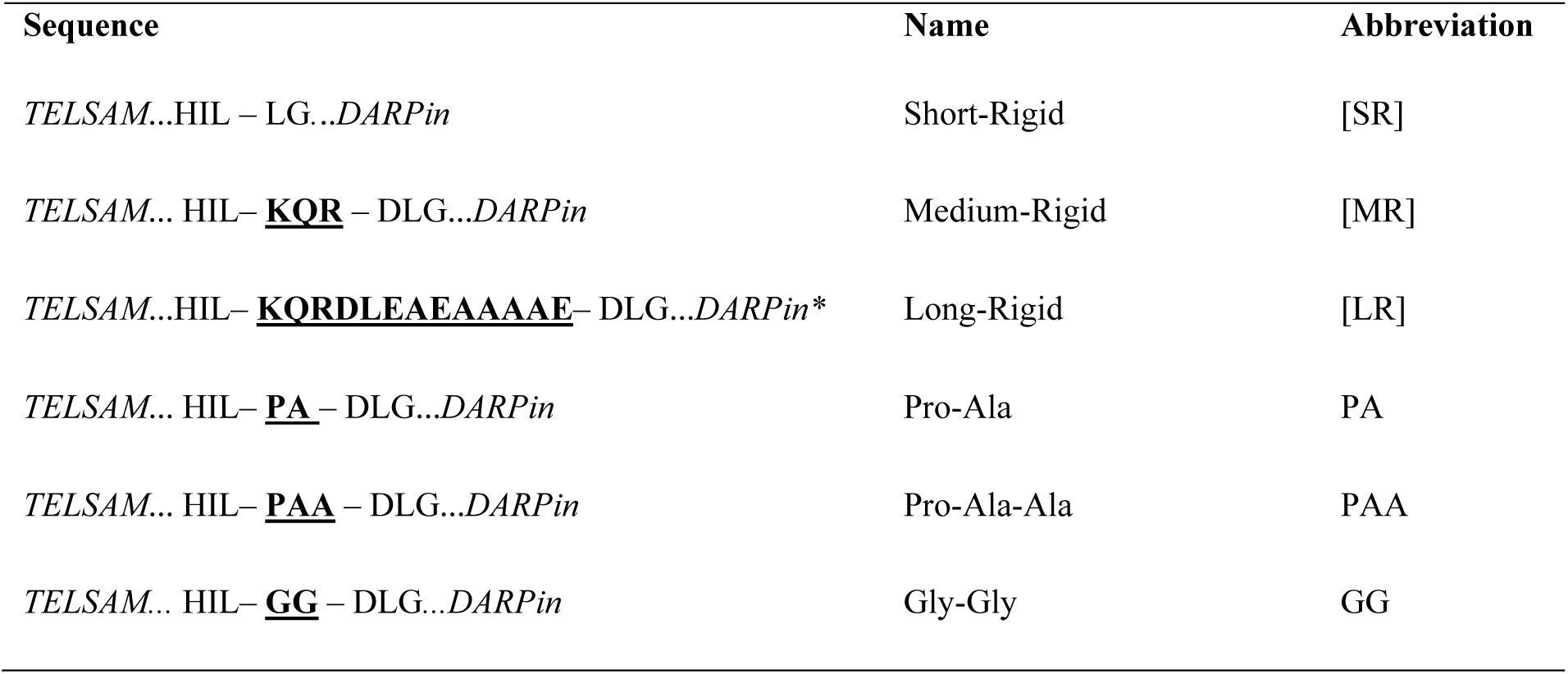
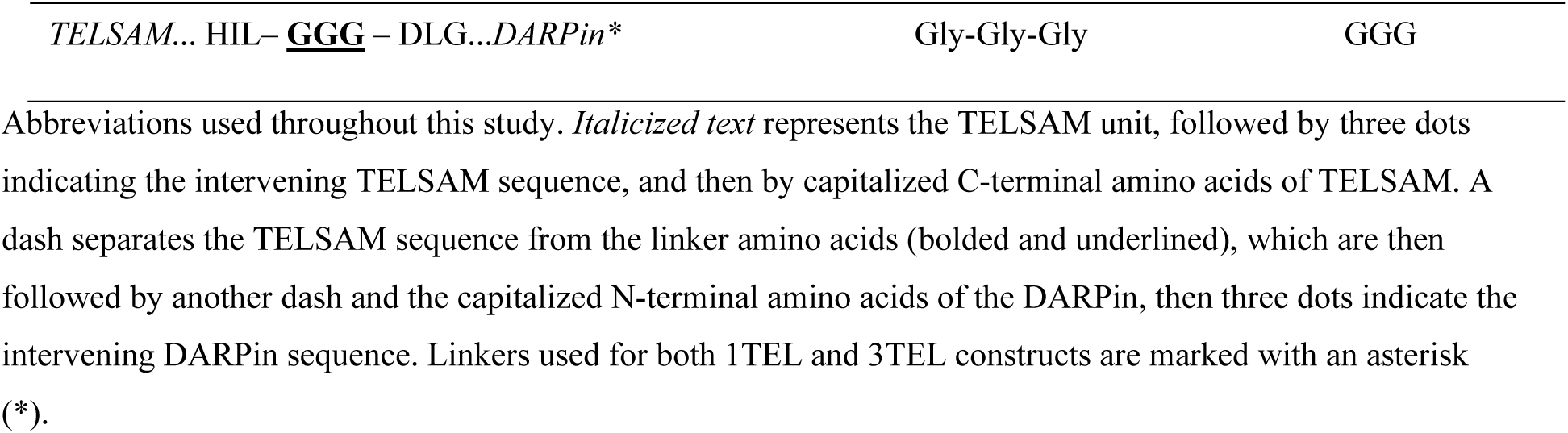
Abbreviations of all Linker Types.

## 2. Results

### 2.1. Steric interference likely prevents 1TEL-GGG-DARPin from forming uniform 1TEL fusion polymers

In our previous experiments, flexible or rigid linkers in 1TEL-DARPin and 3TEL-DARPin constructs were compared. 1TEL constructs consist of a single TELSAM subunit genetically fused to the DARPin while 3TEL constructs consist of three TELSAM subunits fused in tandem and linked by long flexible linkers between their termini (Nauli et al., 2007; Poulos et al., 2017). The DARPin is fused to the C-terminus of the 3^rd^ TELSAM unit. Flexible linkers consisted of Gly-Gly-Gly (GGG) and rigid linkers consisted of a continuous α-helical fusion of the TELSAM C-terminus and the DARPin N-terminus, with 13 strongly helical amino acids (KQRDLEAEAAAAE, as in Nauli et al., 2007) placed between the two (forming a long-rigid linker, [LR]). The four constructs were as follows: 1TEL-GGG-DARPin, 1TEL-[LR]-DARPin, 3TEL-GGG-DARPin, and 3TEL-[LR]-DARPin.

All constructs gave crystals, but only 3TEL constructs, 3TEL-GGG-DARPin (PDB ID: 9E4Q) and 3TEL-[LR]-DARPin (PDB ID: 7N2B (Nawarathnage et al., 2022)), yielded crystals of sufficient quality to provide diffraction data, with resolutions of 3.58 Å and 3.22 Å, respectively. The complete structure of 3TEL-GGG-DARPin is shown in **Figure 1A-C**. The structure shows that the three unique crystal contacts, all of which were DARPin-1TEL contacts, primarily rely on van der Waals interactions with a few hydrogen bonds. This 3TEL-GGG-DARPin structure is remarkable in that, similar to the only other published example of a 3TEL fusion construct, 3TEL-[LR]-DARPin, it exhibits 3TEL polymers arranged in successive layers of polymers, with each layer having an N→C polymer orientation opposite of the layers before and after it. The only other published 3TEL fusion structure is PDB ID 5L0P, which did not exhibit alternating polymer orientations (Poulos et al., 2017).

**Figure 1.**
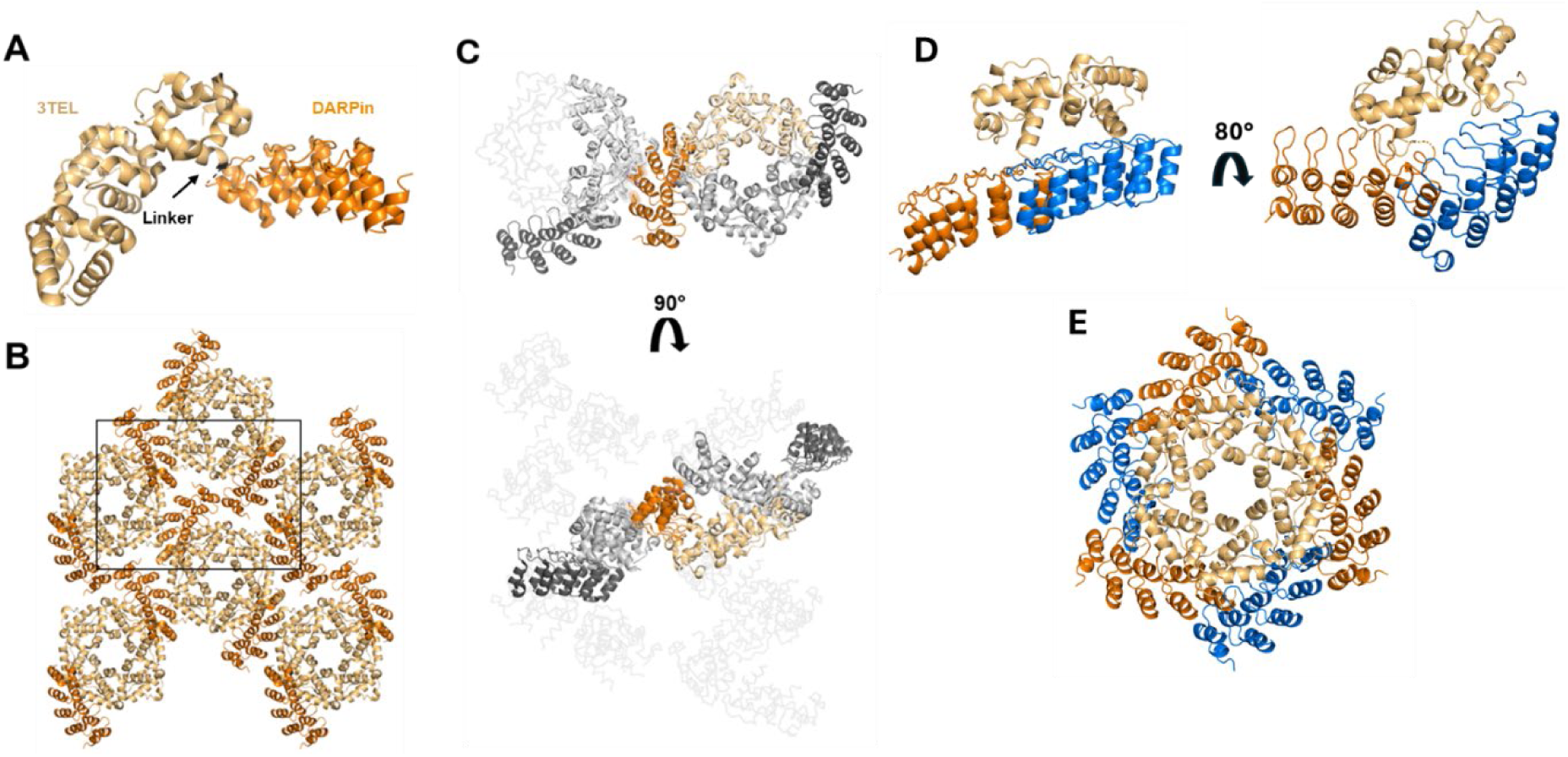
Structural and lattice interactions of the 3TEL-GGG-DARPin. **A.** Structure of the 3TEL-GGG-DARPin monomer (PDB ID: 9DB5), with 3TEL shown in pale orange and DARPin in bright orange. **B.** The 3TEL-GGG-DARPin unit cell (space group P2_1_2_1_2_1_) is outlined in black, with symmetry mates displayed in the same colouring as panel A. **C.** Crystal lattice contacts in this crystal, with 3TEL polymers shown as ribbons and symmetry mates displayed in grey. 3TEL regions are light grey, DARPin regions are dark grey, and the monomer is coloured as in panel A. **D.** The DARPin binding mode from the 3TEL-GGG-DARPin structure superimposed onto successive TELSAM subunits of a 1TEL construct. **E.** As in D, but for one helical turn of the traditional 1TEL P6_5_ polymer.

Since the DARPin in 3TEL-GGG-DARPin adopted a stable binding mode against the polymer and produced reasonable diffraction data, we questioned why 1TEL-GGG-DARPin crystals failed to produce diffraction-quality crystals. To answer this, we superimposed the 3TEL-GGG-DARPin DARPin binding mode onto a 1TEL polymer, as shown in **Figure 1 D-E**. This analysis revealed that the DARPin binding mode observed in the 3TEL-GGG-DARPin structure could not be accommodated if there were six copies of the DARPin per turn of the TELSAM polymer (as in 1TEL-GGG-DARPin). Had the 1TEL-GGG-DARPin adopted this binding mode, the DARPins would clash with each other, preventing them from all adopting the same binding mode against the 1TEL polymer. This situation would prevent uniform polymer-polymer interactions and thus prevent formation of a regular ordered crystallographic lattice, required for high-quality diffraction.

### 2.2. Previous work reveals the use of 10xHis tags can impact the preferred binding mode of the target protein

In our previous 1TEL-CMG2 study (Gajjar et al., 2023), a cleavable 10xHis-SUMO tag was incorporated to enhance protein solubility and facilitate purification. This approach presents a trade-off. During protein purification, the SUMO protease used to cleave the 10xHis tag can result in the proteolytic cleavage of the target protein, which can disrupt TELSAM polymer-polymer association and impair crystallization. However, as observed in our prior study (Gajjar et al., 2023), retention of the His tag can interfere with selected docked orientations of the target protein. We postulate that in some constructs, the His tag can block docking orientations necessary for efficient crystallization.

To further evaluate the impact of the His tag on TELSAM polymerization and crystallization, we generated constructs with and without the His tag. Rigid 1TEL—target linkers remove the rate-limiting step of target protein docking to its host polymer but also limit the conformational flexibility of the target protein, essentially locking it into a single binding mode, irrespective of any His tag. Because of this, for constructs containing rigid linkers (1TEL-[SR]-DARPin, 1TEL-[MR]-DARPin, and 1TEL-[LR]-DARPin), we did not create non-His-tagged versions. In contrast, for constructs with flexible and semi-flexible linkers, which allow the target protein to adopt multiple conformations, we generated both His-tagged and His tag-free versions to evaluate the impact of the His tag on protein behaviour and crystallization.

### 2.3. Crystallization and optimization of 1TEL-DARPin constructs

We tested flexible (GG and GGG), semi-flexible (PA and PAA), and rigid α-helical fusion ([SR], [MR], and [LR]) linkers **(Table 1**). All constructs were expressed, purified, and crystallized independently at least twice by two different teams of students, and all constructs yielded crystals (**Table 1**, **Figure 2**). Overall, rigid linkers crystallized faster than semi-flexible (Pro-Ala_n_) linkers, which in turn crystallized faster than flexible (Gly_n_) linkers, confirming that the rate-limiting step in crystal growth is the docking of the target protein to its host 1TEL polymer. Some constructs gave crystals that were too small for diffraction, and others formed large crystals that nevertheless failed to diffract. This result indicates that crystal size alone is not a predictor of diffraction quality.

**Figure 2.**
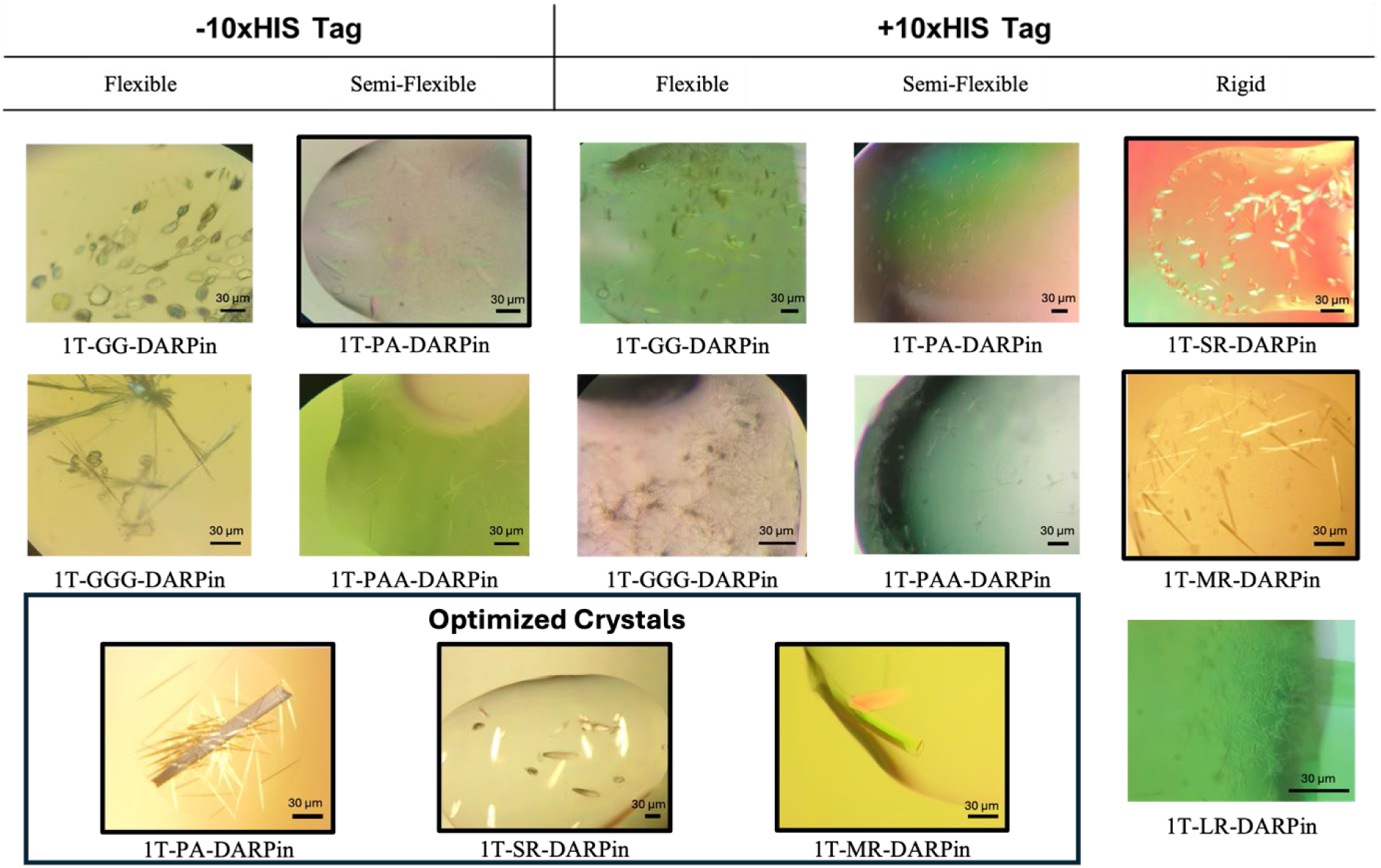
Crystals of 1TEL-DARPin Constructs with Rigid, Semi-flexible, and Flexible Linkers. Representative crystals obtained for all 1TEL-DARPin constructs with rigid, semi-flexible, and flexible linkers, both with (+10xHis Tag) and without (-10xHis Tag) the His-tag. Constructs outlined in black represent those selected for further optimization, resulting in better-quality crystals. The boxed area highlights representative optimized crystals obtained from three constructs: 1T-PA-DARPin, 10xHis-1T-[SR]-DARPin, and 10xHis-1T-[MR]-DARPin.

Among rigid linker constructs, the 10xHis-1TEL-[SR]-DARPin construct crystallized more rapidly than the medium [MR] and long [LR] linker constructs. Additionally, shorter rigid linkers produced larger crystals, suggesting that linker length in helical fusions may influence the quality and/or rate of crystal packing, possibly due to residual motion afforded by the longer α-helical linkers. The short linker may have afforded less residual motion to the DARPin, allowing the 1TEL-DARPin polymers to more quickly dock against each other and thus more readily expand the growing crystal lattice along the a and b axes of the unit cell (**Table 1**).

Most Semi-flexible 10xHis-1TEL-PA- and -PAA-DARPin constructs exhibited poor crystal morphology (pseudo-crystals) and the opposite crystallization propensity trend to that of the rigid linker constructs. While shorter rigid linker constructs produced larger crystals with shorter crystallization times, the semi-flexible linkers Pro-Ala and Pro-Ala-Ala formed equally large crystals, with the Pro-Ala-Ala being 2-8 times faster and diffracting to better resolution but having moderately reduced crystallization propensity. These results show that semi-flexible linker length does not impact crystal size in this 1TEL-DARPin system but does affect crystallization time and crystal quality (**Table 1**). Flexible linkers, GG and GGG, also behaved differently than both rigid and semi-flexible linkers. While flexible 10xHis-1TEL-GG- and -GGG-DARPin constructs exhibited similar crystallization speed and quality trends to the semi-flexible constructs above, the flexible linker constructs exhibited the opposite trend in crystallization propensity and a clear trend in crystal size, with the Gly-Gly linker resulting in larger crystals than the Gly-Gly-Gly linker. In spite of their smaller size, the Gly-Gly-Gly linker crystals were the only flexible-linker crystals to diffract. Crystal morphology was generally poor (needles) across all flexible constructs, which may have impaired their diffraction quality, suggesting that increased target protein conformational freedom may impair the formation of a regular crystal lattice in this 1TEL-DARPin system (**Table 1**).

Removing the 10xHis tag from the semi-flexible and flexible constructs significantly increased their crystallization rate, improved the crystal morphology of the shorter members of each linker type, and markedly improved their crystallization propensity. While removing the 10xHis tag had no significant effect on crystal size, it markedly improved the crystal quality and diffraction limits of the 1TEL-PA-DARPin construct.

Crystals with initial screening diffraction limits better than 4 Å were subjected to crystallization condition optimization. Three constructs met this threshold during screening of sparse conditions: 10xHis-1TEL-[SR]-DARPin, 10xHis-1TEL-[MR]-DARPin, and 1TEL-PA-DARPin. We sought to determine high resolution structures of these three constructs to determine the structural causes of the observed crystallization trends and whether 1TEL-DARPin fusion crystals could surpass 2.0 Å resolution, as observed previously for 1TEL-TNK1.UBA (Nawarathnage et al., 2023) and 10xHis-1TEL-CMG2.vWa (Gajjar et al., 2023). Therefore, we optimized the crystallization conditions of these three constructs by more finely sampling pH and precipitant concentrations in a grid format. **(Table 2).**

**Table 2.** Crystallization time, propensity, and diffraction quality of 1TEL-DARPin constructs in sparse crystallization conditions (before optimization of crystallization conditions)

| Constructs | Days to crystal appearance | Crystal habit | Minor axis of largest crystal in each drop ($\mu\text{m}$) | No. of sparse conditions giving crystals (%) | No. of crystals diffracting X-rays | Best diffraction limit, average, and range ($\text{\AA}$) |
| --- | --- | --- | --- | --- | --- | --- |
| 10xHis- 1T-[SR]-DARPin | 1-2 | Rod | 24-72 | 6.25 | 12 | 3.54 (6.61, 3.54-8.20) |
| 10xHis- 1T-[MR]-DARPin | 0.5-20 | Hexagonal prism | 5-64 | 22.2 | 28 | 3.34 (6.84, 3.34-10.20) |
| 10xHis-1T-[LR]-DARPin | 21-30 | Rod | 10.4 | 2.4 | 0 | No Diffraction |
| 10xHis-1T-[PA]-DARPin | 23-30 | Blob | 32-47 | 1.7 | 0 | No Diffraction |
| 10xHis- 1T-[PAA]-DARPin | 4-9 | Blob | 36-48 | 0.7 | 1 | 4.28 |
| 10xHis-1T-[GG]-DARPin | 21-30 | Needle | 8-40 | 0.7 | 0 | No Diffraction |
| 10xHis-1T-[GGG]-DARPin | 7-14 | Needle | 15-24 | 2.1 | 1 | 12.80 |
| 1T-[PA]-DARPin | 0.5-169 | Rod | 4-89 | 4.9 | 3 | 3.51 (5.00, 3.51-8.10) |
| 1T-[PAA]-DARPin | 1-2 | Needle | 22-43 | 5.2 | 3 | 5.4 (6.9, 5.4-8.2) |
| 1T-[GG]-DARPin | 17-175 | Short barrels | 29-41 | 8.0 | 0 | No Diffraction |
| 1T-[GGG]-DARPin | 5-179 | Needle | 8-16 | 5.9 | ND | Not Harvestable |

### 2.4. Condition optimization did not significantly improve the resolution of 10xHis-1TEL-[MR]-DARPin crystals

The 10xHis-1TEL-[MR]-DARPin originally gave crystals diffracting to an average of 3.34 Å resolution. Following condition optimization, [MR]’s crystals diffracted to an average of 3.28 Å. However, crystals with the best resolution were very mosaic, and did not perform well in data processing, so the structure was solved with a non-optimized crystal that diffracted to 3.47 Å. The resulting structure shows how the rigid medium-length linker influences crystal packing (**Figure 3**). The [MR] linker essentially ‘locks’ the DARPin into one conformation, and while its rigidity stabilizes the protein structure, it results in fewer crystal contacts within the lattice (**Figures 3D-E**, **Table 3**). Analysis of the crystal lattice reveals that weak interactions stabilize the host DARPin, which is consistent with the observed resolution. There are two crystal contacts stabilizing the host DARPin, one forming a charged hydrogen bond between Arg31 of the host DARPin and Asp108 of the 1TEL polymer subunit one helical turn below it (**Figure 3D).** The other crystal contact is a weak hydrogen bond between Asn125 of the host DARPin and the backbone of Glu239 on an adjacent DARPin (**Figure 3E).** This contact is additionally stabilized by van der Waals interactions and has an interface area of 157 Å² **(Figure 3E**). These sparse and relatively weak interactions likely contributed to the limited resolution observed with this construct.

**Figure 3.**
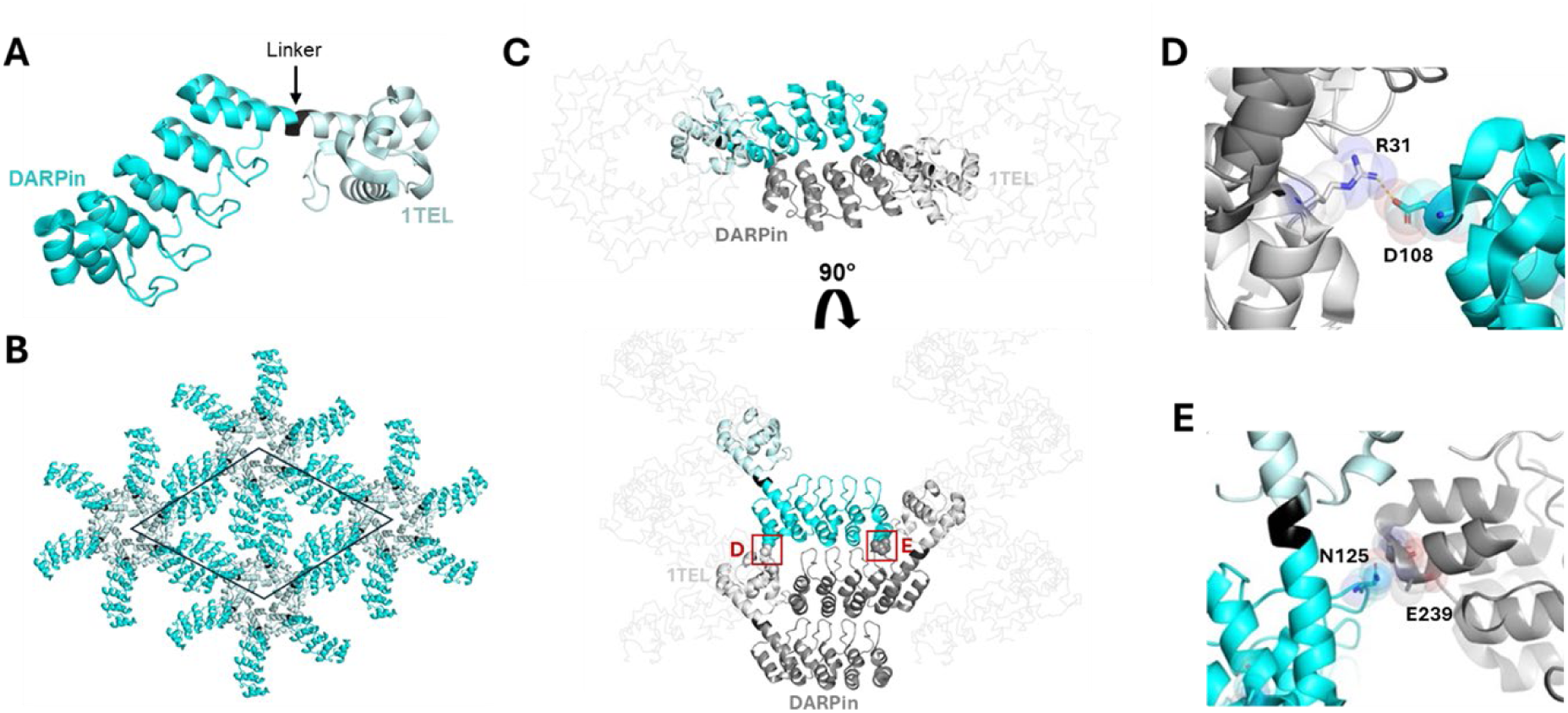
Structure and lattice interactions of a 1TEL-[MR]-DARPin crystal. **A.** Structure of the 10xHis-TEL-[MR]-DARPin monomer (PDB ID: 9DP8), with 1TEL shown in pale cyan, the [MR] linker in black, and DARPin in bright cyan. **B.** The 10xHis-1TEL-[MR]-DARPin unit cell (space group P6_5_) is outlined in black, with symmetry mates displayed in the same colouring as in panel A. **C.** Crystal lattice representation with 1TEL polymers shown as ribbons and symmetry mates displayed in grey. 1TEL regions are light grey, DARPin regions are dark grey, and the monomer is coloured as in panel A. **D.** Close-up view of the interaction between the DARPin (bright cyan) and the 1TEL unit one helical turn below it (light grey). Proteins are depicted in cartoon representation, with interacting residues shown as sticks and spheres. **E.** As in panel D. but showing interactions between the DARPin (bright cyan) and the adjacent DARPin from a neighbouring polymer.

**Table 3.** Diffraction and Data Collection Statistics of optimized crystals.

| Construct | No. of datasets collected | Diffraction limit of collected datasets (Å) | Fraction of non-ice reflections indexed (%) | Average estimated mosaicity (°) | Final ISa of data processing | Unit cell solvent content (%) |
| --- | --- | --- | --- | --- | --- | --- |
| 10xHis-1TEL-[SR]-DARPin (3-Fold) | 3 | 1.85 (1.78-1.92) | 84.9 (82.8-89) | 0.40 (0.35-0.45) | 28 (25-32) | 45 (42-46) |
| 10xHis-1TEL-[SR]-DARPin (2-Fold) | 2 | 3.53 (3.51-3.54) | 63.5 (53.5-73.5) | 0.35 (0.34-0.35) | 23 (23-24) | 56 (53- 59) |
| 10xHis-1TEL-[MR]-DARPin | 6 | 3.28 (2.87-4.61) | 15.0 (13.0-42.5) | 0.70 (0.40-1.60) | 2 (10-40) | 58.943 (56-60) |
| SUMO 1TEL-[PA]-DARPin | 9 | 2.69 (1.57-4.00) | 51.2 (4.3-93.9) | 0.39 (0.16-0.70) | 22 (11-36) | 54 (50-61) |
| SUMO-1TEL-[GG]-UBA | 9 | 2.01 (1.42 – 4.0) | 31.6 (21.6–44.3) | 0.55 (0.25-1.2) | 23 (16-29) | 44 (41-45) |

### 2.5. Optimized 10xHis-1TEL-[SR]-DARPin crystals exhibit two distinct crystal forms

We observed two distinct crystal forms from the 10xHis-1TEL-[SR]-DARPin construct, to which we will refer to hereafter as the ‘2-fold’ and ‘3-fold’ crystal forms, respectively. The ‘2-fold’ form showed crystal contacts similar to those observed in the 10xHis-1TEL-[MR]-DARPin construct, while the ‘3-fold’ lattice revealed a novel arrangement where DARPins came together in a staircase-like formation along the P3_2_ screw axes of the P6_5_ unit cell. When viewed along the c-axis of the unit cell, three DARPins meet in a triangular arrangement. The ‘3-fold’ lattice featured more crystal contacts, showed higher internal order, and achiever a higher resolution compared to the ‘2-fold’ lattice.

#### 2.5.1. 10xHis-1TEL-[SR]-DARPin ‘2-Fold’ crystal form resembles the [MR] crystal form, with relatively weak contacts

We solved the structure of a ‘2-fold’ 10xHis-1TEL-[SR]-DARPin crystal at 3.54 Å resolution. The structure exhibits a similar lattice architecture to the 10xHis-1TEL-[MR]-DARPin construct (**Figure 3A**), though with distinct crystal contacts. Analysis of this structure revealed three key interaction points stabilizing the crystal lattice. The first was a DARPin-DARPin crystal contact stabilized by a hydrogen bond. (**Figure 4D**). The remaining two contacts were DARPin-1TEL interactions. One includes van der Waals interactions and a salt bridge between the host DARPin and the 1TEL unit one helical turn below it in the same 1TEL polymer (**Figure 4E**). The last contact is another weak van der Waals interaction between the host DARPin Ile228 and E233 and Y26 of the 1TEL subunit of a neighbouring 1TEL polymer **(Figure 4F**). The total interface area was 307 Å^2^. The weaker nature of these limited interactions also likely contributed to the lower resolution achieved in this construct.

**Figure 4.**
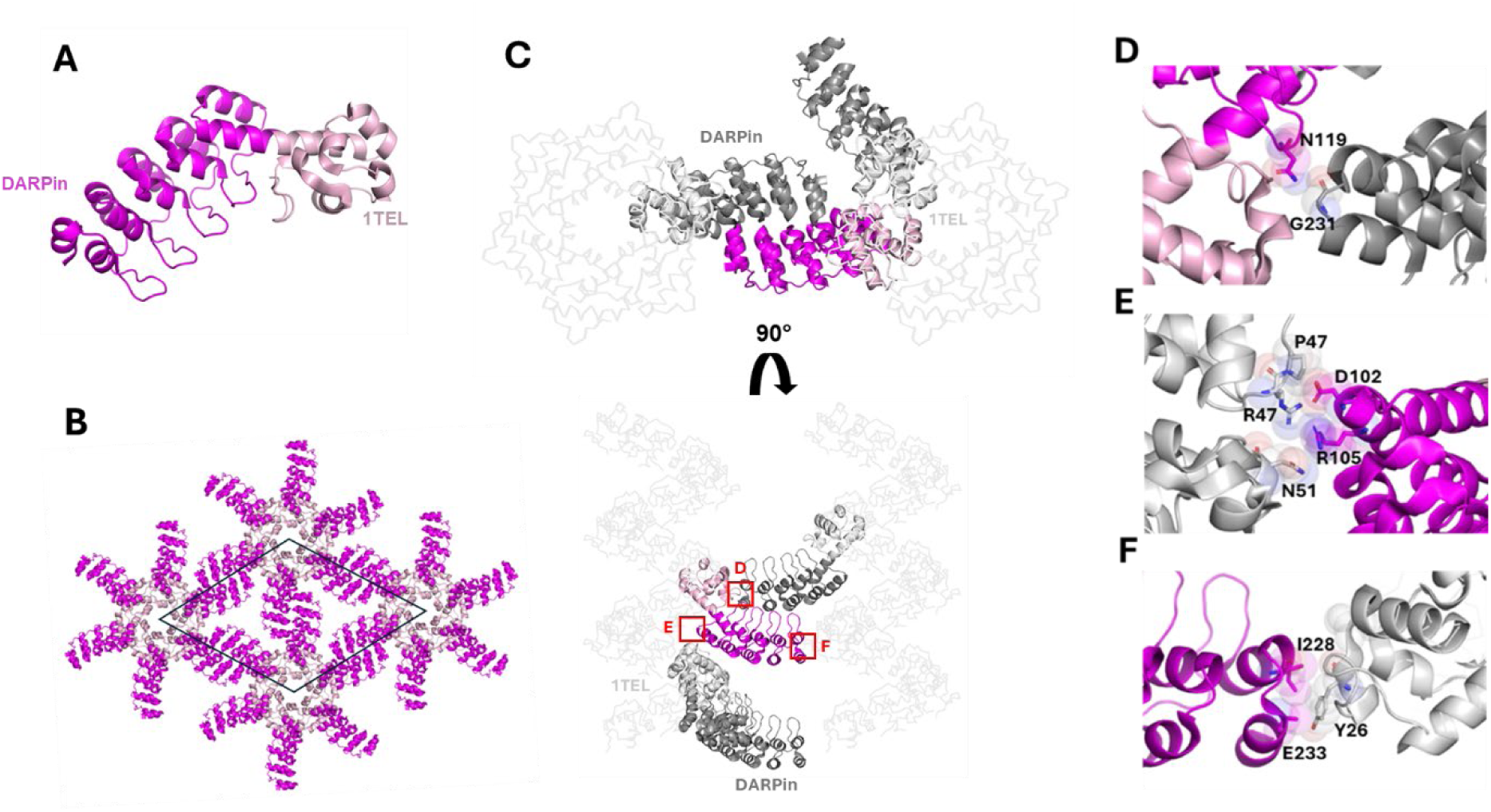
Structure and lattice interactions of a ‘2-fold’ 1TEL-[SR]-DARPin crystal. **A.** Structure of the 2-fold 1TEL-[SR]-DARPin monomer (PDB ID: 9DVG), with 1TEL shown in pale pink and DARPin in magenta. **B.** The 1TEL-[SR]-DARPin unit cell (space group P6_5_) is outlined in black, with symmetry mates displayed in the same colouring as in panel A. **C.** Crystal lattice representation with 1TEL polymers shown as ribbons and symmetry mates displayed in grey. 1TEL regions are light grey, DARPin regions are dark grey, and the monomer is coloured as in panel A. **D.** Close-up view of the interaction between two DARPins, with interacting residues shown as sticks and spheres. **E-F.** Two unique interactions between the DARPin (magenta) and nearby 1TELs (light grey) from the host polymer or a neighbouring polymer.

#### 2.5.2. The ‘3-fold’ crystal form of 10xHis-1TEL-[SR]-DARPin exhibits enhanced crystal contacts

We solved the structure of a ‘3-fold’ 10xHis-1TEL-[SR]-DARPin crystal at 1.78 Å resolution. In this structure we found a unique staircase-like structure along the P3_2_ axes of the unit cell, where DARPins assembled in a 3-fold helical arrangement. This crystal achieved a better resolution likely due to enhanced crystal contacts. Unlike the ‘2-fold’ crystal form, the ‘3-fold’ crystal form featured two unique DARPin-DARPin crystal contacts per asymmetric unit, each involving more residues and stronger interactions, including some salt bridges. The first contact involved hydrogen bonds between two side chains of the host DARPin and the backbones of two lysine residues of a neighbouring DARPin (**Figure 5D**). The second contact occurred between the host DARPin and a neighbouring DARPin subunit and included a salt bridge and a hydrogen bond stabilized by van der Waals interactions (**Figure 5E**). The final interaction consisted of two hydrogen bonds between the host DARPin and a neighbouring 1TEL subunit from another 1TEL polymer (**Figure 5F)**. The total interface area of all three interactions was 1085 Å^2^. These extensive interactions are likely responsible for the increased internal order and higher resolution observed with this construct.

**Figure 5.**
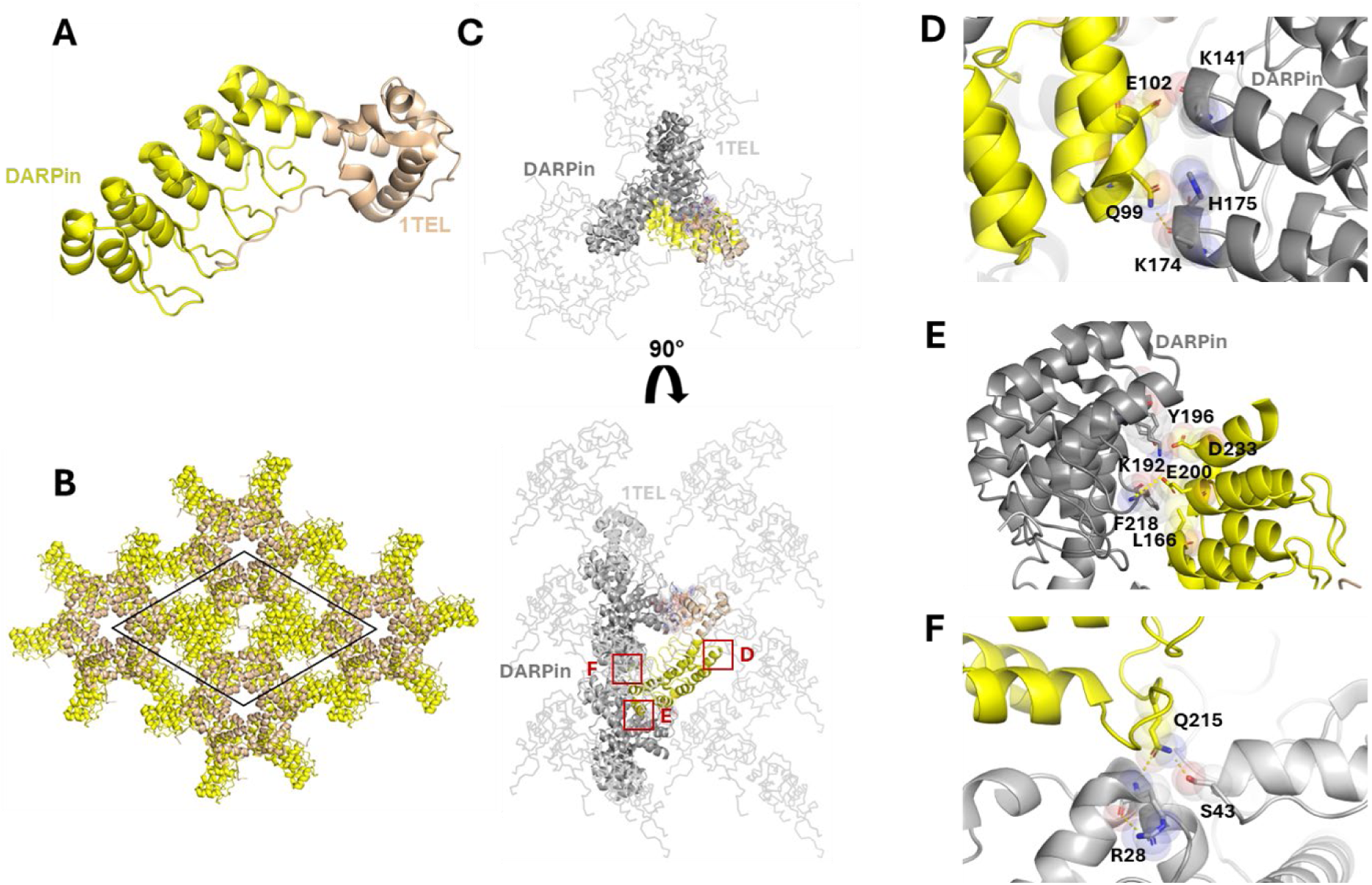
Structure and lattice interactions of a ‘3-fold’ 1TEL-[SR]-DARPin crystal. **A.** Structure of the ‘3-fold’ 10xHis-1TEL-[SR]-DARPin monomer (PDB ID: 9DAM), with 1TEL shown in wheat and DARPin in yellow. **B.** The 3-fold 10xHis-1TEL-[SR]-DARPin unit cell (space group P6_5_) is outlined in black, with symmetry mates displayed in the same colouring as in panel A. **C.** Crystal lattice representation with 1TEL polymers shown as ribbons and symmetry mates displayed in grey. 1TEL regions are light grey, DARPin regions are dark grey, and the monomer is coloured as in panel A. **D-E.** Close-up view of the interactions between the DARPin (yellow) and the adjacent DARPin from a neighbouring polymer (dark grey). Proteins are depicted in cartoon representation, with interacting residues shown as sticks and spheres. **F.** As in panels D-E but showing interactions between the DARPin (yellow) and a 1TEL subunit (light grey) one helical turn above it in the same 1TEL polymer. Also shown are salt bridge contacts, represented by a yellow dotted line.

### 2.6. 1TEL-PA-DARPin achieved the highest resolution of all constructs and exhibited the most crystal contacts

The 1TEL-PA-DARPin construct produced the highest-resolution structure in this study at 1.57 Å, slightly better than the resolution of this same DARPin crystallized on its own (wwPDB ID: 4J7W, Seegar et al) and exhibited strong crystal contacts. We observed rapid crystal dissolution during harvesting but were able to harvest several crystals quickly enough to outpace complete dissolution. Unlike the rigid linker constructs, this construct lacked a 10xHis tag. The DARPin was observed to form four unique crystal contacts per asymmetric unit. The first involved van der Waals contacts between side chains of the host DARPin and various side chains of a neighbouring DARPin, mediated by acetate ions (**Figure 6D**). The second contact occurred between the long finger loops connecting the α-helices of the DARPin and an adjacent 1TEL and consisted largely of hydrogen bonds, some of them water mediated, and a van der Waals contact, with the entire interaction having an area of 470 Å² (**Figure 6E**). The third contact was another DARPin-DARPin van der Waals interaction (**Figure 6F**) with an interface area of 420 Å². The final contact involved two hydrogen bonds between Lys134 of the focus DARPin and residues His74, Pro77, and Ala78 of a neighbouring 1TEL (**Figure 6G)**. The total interface area was 1855 Å^2^, the greatest observed among all the constructs.

**Figure 6.**
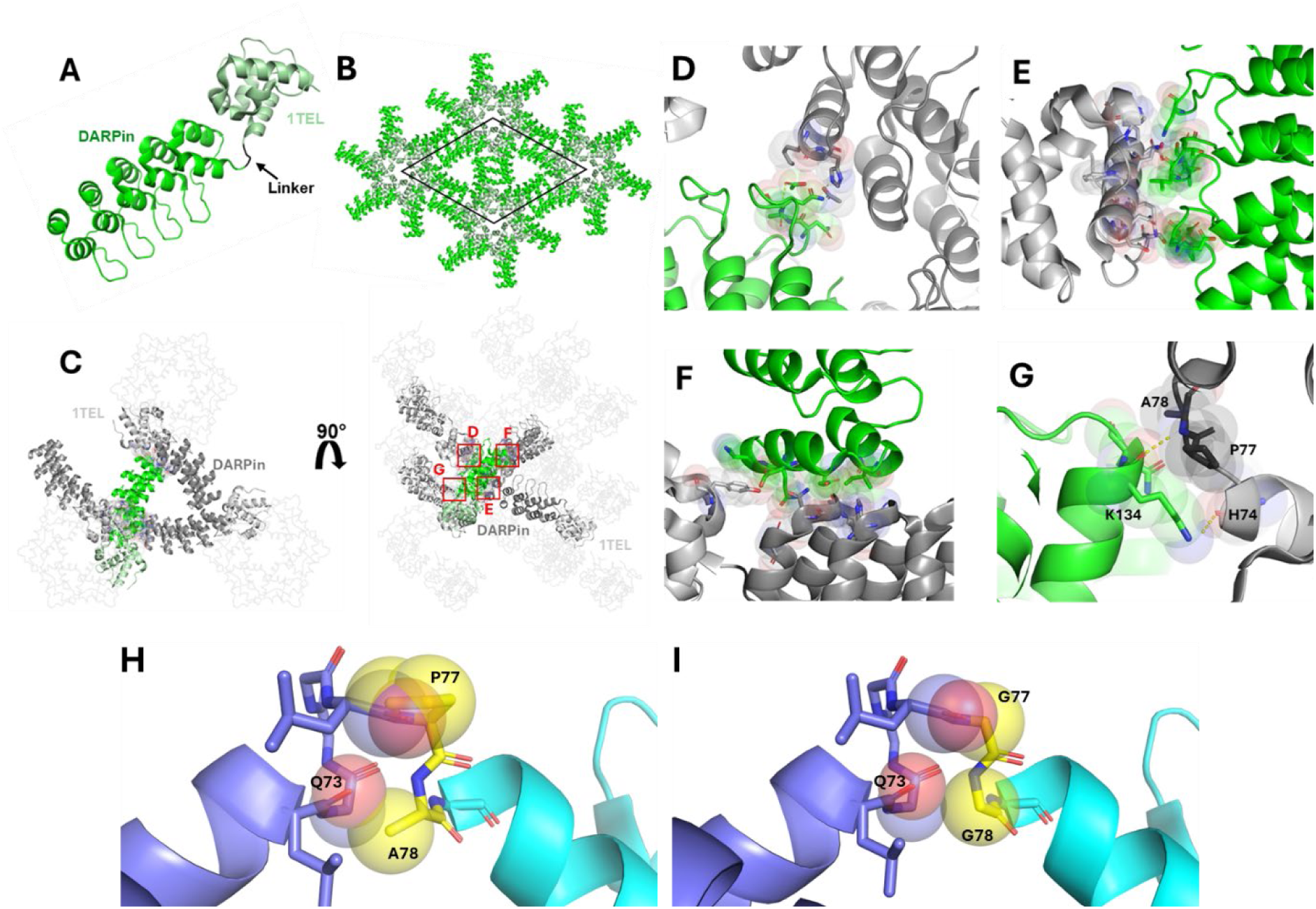
Structure and lattice interactions of a 1TEL-[PA]-DARPin crystal. **A.** Structure of the 1TEL-[PA]-DARPin monomer (PDB ID: 9DB5), with 1TEL shown in pale green and the DARPin in bright green. **B.** The 1TEL-[PA]-DARPin unit cell (space group P65) is outlined in black, with symmetry mates displayed in the same colouring as in panel A. **C.** Crystal lattice representation with 1TEL polymers shown as ribbons and symmetry mates displayed in grey. 1TEL regions are light grey, DARPin regions are dark grey, and the monomer is coloured as in panel A. **D-G**. Close-up views of the interactions between the host DARPin (bright green) and adjacent DARPin proteins from neighbouring polymers or neighbouring 1TEL units (light grey). Proteins are depicted in cartoon representation, with interacting residues shown as sticks and spheres and hydrogen bonds shown as yellow dotted lines. **H**. Superposition of a theoretical Pro-Ala linker between a 1TEL and a UBA (cyan) and the published structure of 1TEL-GG-UBA (PDB ID: 7TDY) with clashes shown in spheres. **I.** The published 7TDY structure with its original Gly-Gly linker.

#### 2.6.1. S-1T-PA-DARPin crystals likely dissolved due to a pH shift caused by acetic acid evaporation

During crystal harvesting, we observed that 1TEL-PA-DARPin crystals immediately began dissolving once the well was opened. From the structure, we noted that many crystal contacts consisted of side-chain carboxylic acids hydrogen bonding each other. While direct carboxylic acid-carboxylic acid hydrogen bonds are usual in biological systems, they were possible in this pH 4.6 crystallization drop as a significant number of these carboxylates (pKas around pH 4) would be protonated at pH 4.6. The juxtaposition of these carboxylic acids likely also raised their pKas above 4. Numerous acetates were also visible in the electron density. Upon opening the well for crystal harvesting, acetic acid molecules undoubtedly began evaporating, shifting the equilibrium of the acetates remaining in solution toward the protonated state, and raising the pH of crystal drop. This increase in pH likely began deprotonating the carboxylic acid side chains, causing them to repel each other and disrupt the corresponding crystal contacts, resulting in rapid crystal dissolution. As few TELSAM fusion crystals appear in crystallization conditions containing volatile buffer components and even fewer involve carboxylic acid-carboxylic acid hydrogen bonds, we expect that this phenomenon will not be typical of TELSAM fusion crystals.

## 3. Generalizability of Linker Optimization to TELSAM-UBA Crystallization

After determining that the semi-flexible Pro-Ala linker was most effective for the 1TEL-DARPin system, we sought to validate this result against a second protein target. We selected the ubiquitin-associated (UBA) domain of human thirty-eight negative kinase 1 (TNK1) as we have previously characterized the 1TEL-UBA fusion and because the UBA domain also has an α-helical N-terminus. Since the [SR] linker also produced 1TEL-DARPin crystals that diffracted to an acceptable resolution, we decided to test it alongside the Pro-Ala linker. We previously reported the structure of this same UBA domain at a resolution of 1.53 Å using 1TEL fusion with a 2x-Gly linker (Nawarathnage et al., 2023)

We thus tested linking the UBA domain to 1TEL with three distinct linkers: Gly-Gly (GG, positive control), a direct helical fusion [SR], and Pro-Ala (PA). All constructs successfully yielded crystals (**Table 4**), but only 1TEL-GG-TNK1.UBA crystals diffracted X-rays, while crystals of constructs utilizing the [SR] and PA linkers did not diffract. The PA linker performed poorly in the 1TEL-UBA system, apparently due to poor solubility. Rapid precipitation in crystal trays was observed and may have outpaced the formation of regularly aligned TELSAM polymers. This linker-induced solubility reduction warrants future investigation. While the crystals of the ‘PA’ construct were of adequate size and exhibited a hexagonal prism crystal habit, crystals of the ‘[SR]’ construct were too small for diffraction. This indicated that the Gly-Gly linker remained the current best choice for 1TEL-UBA fusion crystallization. The detailed diffraction statistics for this construct are given in **Table 2**.

**Table 4.** Comparison of Unique Crystal Contacts and Total Contact Area Per Asymmetric Unit Across 1TEL-DARPin Constructs.

| Construct | No. of Unique Crystal Contacts per AU involving the DARPin | Total Area of Crystal Contacts (Å<sup>2</sup>) | Percent Solvent Content | Resolution (Å) |
| --- | --- | --- | --- | --- |
| 10xHis-1TEL-[MR]-DARPin | 2 | 199 Å<sup>2</sup> | 59 | 3.47 |
| 10xHis-1TEL-[SR]-DARPin (2-Fold) | 3 | 307 Å<sup>2</sup> | 53 | 3.54 |
| 10xHis-1TEL-[SR]-DARPin (3-Fold) | 3 | 1085 Å<sup>2</sup> | 42 | 1.78 |
| 1TEL-PA-DARPin | 4 | 1855 Å<sup>2</sup> | 50 | 1.57 |

**Table 5.** Crystallization time, propensity, and diffraction quality of 1TEL-UBA constructs.

| Construct | Days to crystal appearance | Crystal minor axis ($\mu\text{m}$) | Fraction of sparse conditions giving crystals (%) | No. of crystals that diffracted > 2 Å | Best diffraction limit (Å) |
| --- | --- | --- | --- | --- | --- |
| 1TEL-GG-UBA | 1 | 80 | 3.5 | 10 | 1.4 |
| 1TEL-PA-UBA | 1 | 30 | 10 | 0 | No Diffraction |
| 10xHis-1TEL-[SR]-UBA | 1 | 12 | 3.8 | 0 | Not Harvestable |

## 4. Discussion

The choice of linker between 1TEL and the target protein appears to be dependent on the protein itself, in a non-intuitive manner. The Gly-Gly linker outperformed the [SR] and Pro-Ala linkers in the context of the 1TEL-UBA system, in stark contrast to the 1TEL-DARPin system, where Pro-Ala_n_ and [SR] linkers performed better than Gly_n_ linkers. Among all the linkers tested in the 1TEL-DARPin system, the semi-flexible PA linker produced the highest resolution and the most extensive crystal contacts. We propose that this differential behaviour is due to the differing sizes of the UBA (9 kDa) and DARPin (17 kDa) target proteins. While the UBA could fold back to lie flat against its host polymer, the DARPin was too large to do so (**Figure 1D, E**). As Gly_n_ linkers appear to work best by enabling the target protein to fold back against their host polymers (as exhibited by the 1TEL-GG-UBA and 3TEL-GGG-DARPin crystal structures), this observation may explain why the Gly_n_ linkers performed poorly in the context of a 1TEL-DARPin fusion. The Pro-Ala linker may have been inflexible enough to allow the DARPin binding mode observed in the 1TEL-PA-DARPin crystal structure but not so rigid as to incur the challenges observed with the rigid α-helical 1TEL-DARPin fusions described in this study. Modelling a Pro-Ala linker in place of the Gly-Gly linker in our published 1TEL-GG-UBA structure (wwPDB ID: 7TDY) (Nawarathnage et al., 2023) reveals that a proline cannot access the Ramachandran angle of the first Gly76 of the Gly-Gly linker without incurring serious steric clashes with the backbone carbonyl of the Leu75 preceding it. Likewise, substituting an Ala for the second Gly77 in this linker results in serious clashes with the backbone carbonyl of Leu72 one helical turn below Gly77 in the same α-helix of the 1TEL domain. In fact, no other amino acids except Gly can be substituted at either of these positions in the solved 1TEL-GG-UBA structure. As the amino acids to before and after the Gly-Gly linker are part of α-helices, it is unlikely that a Pro could have been accommodated at any other position near the 1TEL-UBA junction point (**Figure 6H-I**). When designing this fusion, we anticipated the Pro-Ala linker to force the UBA domain to adopt a new and unique docked orientation relative to the UBA. Since the 1TEL-PA-UBA construct did form crystals, the UBA likely did adopt a new docked orientation, but possibly one not compatible with sufficiently productive crystal contacts.

We also observed that the 10xHis-1TEL-[MR]-DARPin and the ‘2-fold’ 10xHis-1TEL-[SR]-DARPin crystals exhibited similar DARPin orientations, resulting in fewer and weaker crystal contacts and likely contributing to the observed poor resolution. In contrast, the ‘3-Fold’ 10xHis-1TEL-[SR]-DARPin crystal form showed improved crystal contacts and better resolution. These findings suggest that while rigid linkers may reduce the entropic cost of crystal formation by essentially ‘locking’ target proteins in place, they struggle to produce high-quality structures. By restricting the conformational flexibility of the DARPin, rigid linkers prevented the DARPin from sampling alternative binding modes against the 1TEL polymer. While rigid linkers eliminate the rate-limiting target protein docking step otherwise required for polymer-polymer association and crystal growth, they may also hinder the formation of optimal crystal contacts, contributing to the formation of lower-quality crystals and reduced resolution. We hypothesize that the target protein binding mode (to its host TELSAM polymer) observed in solved structures using flexible and semi-flexible linkers is not necessarily the lowest-energy binding mode, but rather a binding mode that enables productive polymer-polymer contacts. These crystal contacts then contribute to improved internal order within the crystal, leading to higher resolution. Rigidly fused target proteins have no opportunity to sample multiple binding modes in search of a binding mode that gives productive crystal contacts and so exhibit a lower (but non-zero) probability of forming strong polymer-polymer contacts. In support of this hypothesis, crystals of the 10xHis-1TEL-[MR]-DARPin construct in this study exhibited poor diffraction limits even after crystallization condition optimization. Likewise, a 1TEL-[LR]-UBA fusion did not form crystals while a 1TEL-[SR]-Claudin1-variant fusion formed small needles that failed to diffract X-rays (suggesting a defect in polymer-polymer association), and our previous 3TEL-[LR]-DARPin fusion formed only thin plates, exhibiting a clear defect in target protein-target protein crystal contacts. Additionally, in the hands of different teams of students, both a 1TEL-[SR]-UBA fusion (identical to the one in this study) and a 1TEL-[SR]-Vacuolar Protein Sorting-34 fusion only crystallized in non-polymer crystal forms, to be described elsewhere.Of note is the fact that the position of the DARPin domain between the ‘3-fold’ and ‘2-fold’ 1TEL-[SR]-DARPin structures differs by as much as 19.3 Å at the distal end of the DARPin, due to some flexibility in the conformation of the helical linker (specifically residues Leu87-Leu88). This observation confirms that even ‘rigid’ α-helical fusions retain some degree of flexibility, as previously posited (Nawarathnage et al., 2022) (**Figure 8A**). This subtle displacement of the DARPin likely contributed to the differing crystal packing between the 2-fold’ and ‘3-fold’ 1TEL-[SR]-DARPin structures. Superpositions of rigid linkers (**Figure 8A)**, flexible and semi-flexible linkers (**Figure 8B)** and all DARPin constructs together (**Figure 8C)** highlight the impact of linker composition on DARPin orientation, docking, crystal contacts, and overall crystal formation.

In this study, the semi-flexible PA linker produced the highest resolution and most ordered crystals for the 1TEL-DARPin system. However, the Gly-Gly linker remained the most effective for the 1TEL-UBA system, confirming that optimal linker choice is dependent on the target protein. These findings support selecting a linker that provides enough flexibility for the target protein to adopt a docking conformation to its host polymer capable of supporting productive inter-polymer contacts. We thus recommend prioritizing short flexible or semi-flexible linkers, such as Gly-Gly or Pro-Ala, over rigid linkers, to maximize the odds of productive crystal contact formation. We further recommend that rigid linkers (due to their tendency to limit crystal quality and diffraction resolution) be reserved for cases where more flexible linkers have failed. Finally, we recommend avoiding the retention of N-terminal poly-histidine tags for flexibly or semi-flexibly-fused constructs.

**Figure 7.**
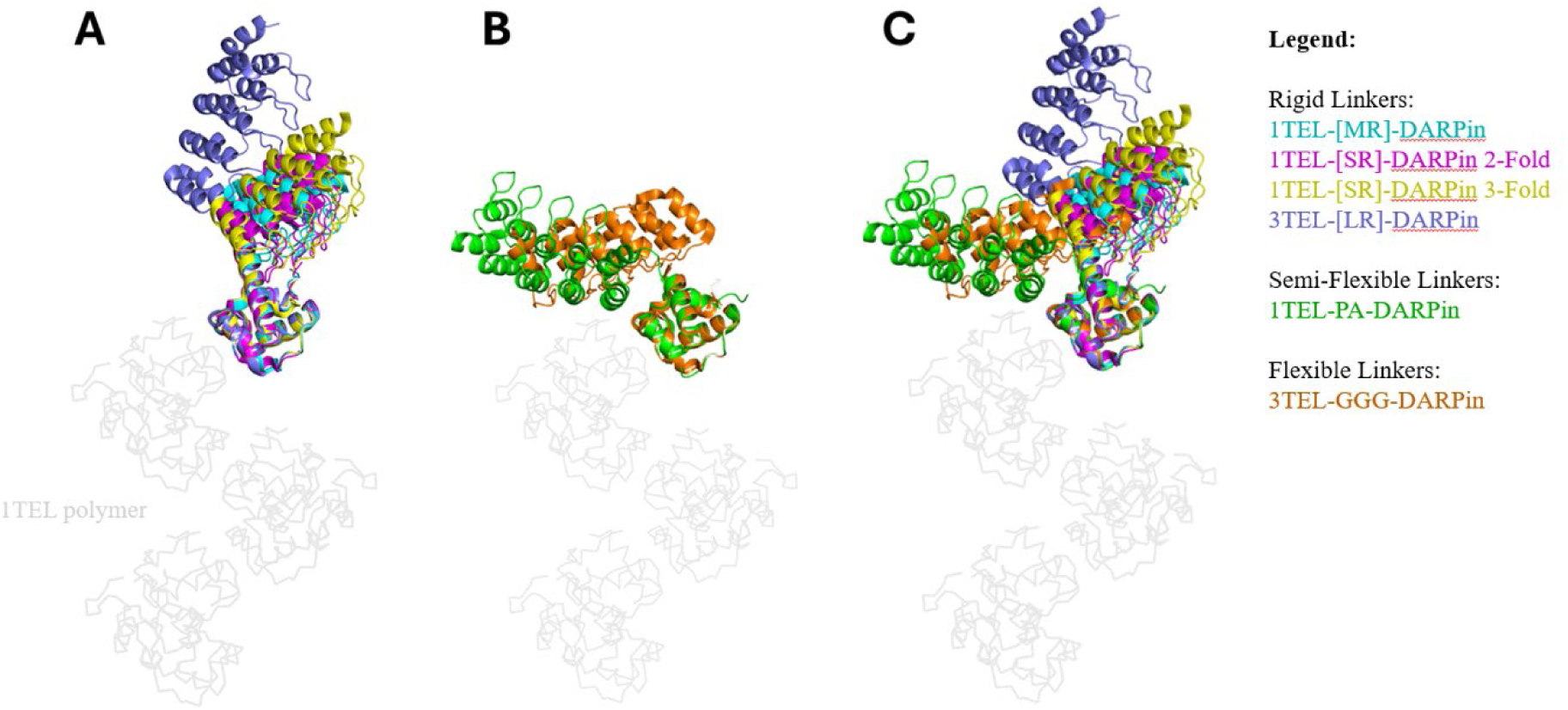
Superposition of rigid, semi-flexible, and flexible TELSAM-DARPin fusions. **A.** Structures of constructs with rigid linkers: 1TEL-MR-DARPin (PDB ID: 9DP8, cyan), 1TEL-[SR]-DARPin 3-fold (PDB ID: 9DAM, yellow), 1TEL-[SR]-DARPin 2-fold (PDB ID: 9DVG, pink), and 3TEL-[LR]-DARPin (PDB ID: 7N2B, blue), superimposed onto a 1TEL polymer (light grey). **B.** Structures of constructs with semi-flexible and flexible linkers, 1TEL-PA-DARPin (PDB ID: 9DB5, green) and 3TEL-GGG-DARPin (PDB ID: 9E4Q, orange), superimposed onto a 1TEL polymer (light grey). **C.** Superposition of A and B.

## 5. Materials and Methods

### 5.1. Cloning

A gene fragment that placed the sequence of the Designed Ankyrin Repeat Protein (DARPin) from PDB ID 4J7W (Seeger et al., 2013) was placed downstream of residues 47-123 (the SAM domain) of the human translocation ETS leukaemia protein, (TEL, Uniprot: P41212, PDB ID 2QAR). This variant of the SAM domain bears a V112E mutation (that makes its polymerization pH-sensitive) and will be referred to as 1TEL hereafter. To form the flexible and semi-flexible linkers, Pro-Ala, Gly-Gly, Pro-Ala-Ala, and Gly-Gly-Gly were inserted between the 1TEL and DARPin domains. To form the short rigid [SR] linker, the first amino acid of the DARPin was deleted and the 1TEL and DARPin sequence fragments were directly fused. To form the medium rigid [MR] linker, the amino acids Lys-Gln-Arg were inserted between the 1TEL and DARPin domains using Geneious (Geneious 9.1.8, https://www.geneious.com). To form the long helical [LR] linker, the helix-forming sequence Lys-Gln-Arg-Asp-Leu-Glu-Ala-Glu-Ala-Ala-Ala-Ala-Glu was inserted between the 1TEL and DARPin domains. No mutations were introduced into the DARPin, except the single amino acid N-terminal truncation to shorten the α-helical linker between 1TEL and the DARPin in the 10xHis-1TEL-[SR]-DARPin construct.

Similarly, Fusions of 1TEL with the human Thirty-eight negative Kinase-1 (TNK1) ubiquitin-associated (UBA) domain (residues 590-666, Uniprot Q13470) were designed using Foldit (Kleffner et al., 2017) and PyMol (https://pymol.org/2/,Schrödinger). We substituted Pro-Ala (semi-flexible) or a single Lys (rigid, [SR]) in place of the Gly-Gly linker of the 1TEL-GG-UBA construct from wwPDB ID: 7TDY (Nawarathnage et al., 2023). In the case of the 10xHis-1TEL-[SR]-UBA fusion, Leu591 of the UBA domain also had to be mutated to Val to resolve a steric clash. The 1TEL-GG-UBA and 1TEL-PA-UBA constructs were placed downstream of a 10xHis-SUMO domain to allow for His tag removal prior to crystallization, while the 10xHis-1TEL-[SR]-UBA construct was cloned immediately downstream of a permanent 10xHis tag.

Gene fragments were synthesized by Twist Bioscience (https://www.twistbioscience.com) and were assembled into the pET42_SUMO vector after cutting with XhoI (GG_n_ and PA_n_ constructs) or with XhoI and NdeI ([SR], [MR], and [LR] constructs) (Walls et al., 2022) using Gibson assembly (Gibson et al., 2009). The plasmids were then introduced into BL21(DE3) cells and sequence verified using Sanger sequencing in both directions by Eton Bioscience (https://www.etonbio.com)

### 5.2. Protein Expression

Lysogeny Broth (Luria-Bertani, LB) medium supplemented with 100 µg/ml kanamycin and 0.35% glucose was inoculated with colonies from transformation or 10 µL frozen cell stock and shaken at 37 °C and 250 rpm overnight for approximately 16 hours. The next day, 10 mL of the overnight culture were added to 1 L LB media supplemented with 0.05% glucose and 50 ug/mL kanamycin and incubated at 37 °C and 250 rpm. The optical density (OD) was measured every 30 minutes until it reached 0.8, at which point 100 µL of 1 M isopropyl β-D-1-thiogalactopyranoside (IPTG) was added. The culture was then incubated at 18°C and 250 rpm overnight. The following day, the cells were collected by centrifugation, snap-frozen in liquid nitrogen, and stored at −80°C.

### 5.3. Purification of 1T-GG-DARPin, 1T-PA-DARPin, 1T-GGG-DARPin, 1T-PAA-DARPin, 1T-GG-UBA, 1T-PA-UBA, and 3TEL-GGG-DARPin (with SUMO cleavable tag)

All purification steps were performed on ice or in a 4°C refrigerator. Cell pellet was weighed and lysed using 3 mL of wash buffer (50 mM Tris, pH 8.8, 200 mM KCl, 50 mM imidazole) per gram of wet cell paste supplemented with 0.3 mg/mL lysozyme, 0.03 mg/mL deoxyribonuclease I, 0.2 mg/mL Phenylmethylsulfonyl-fluoride (PMSF), and 1% methanol. The lysate was run through a homogenizer twice with a nozzle pressure of 18,000 psi (NanoDeBEE 45-2, Pion, Inc.). During one purification of SUMO-1TEL-PA-DARPin, cell lysis was instead accomplished via sonication at 60% power with 12 s on/59 s off for 25 cycles (Qsonica Q500) in a spinning ice bath. The resulting lysate was centrifuged at 40,000 g and the supernatant was loaded onto 2 mL of HisPure Ni-NTA resin (Thermo Scientific). The column was then washed with 10 column bed volumes (CV) of wash buffer (50 mM Tris, pH 8.8, 200 mM KCl, 50 mM imidazole). The column was eluted with 10 mL of elution buffer (50 mM Tris, pH 8.8, 200 mM KCl, 400 mM imidazole) until protein was no longer detected using Bradford reagent (Bradford, 1976). The elute fraction was then desalted using several PD-10 desalting columns (Cytiva) in parallel with 2.5 mL of elution fraction added to each desalting column. The SUMO tag was removed by incubating the protein overnight at 4°C with 0.5 mg SUMO protease per 100mL of protein (Lau et al., 2018). The SUMO protease and cleaved SUMO tags were removed by flowing the cleavage reaction over 2 ml of fresh Ni-NTA resin. The protein was then concentrated to 3 mL and loaded onto a 100 ml Superdex 200 Prep Grade size exclusion column (Cytiva). The final purified proteins were judged to be greater than 95% pure by SDS-PAGE.

### 5.4. Purification of 10xHis-1TEL-GG-DARPin, 10xHis-1TEL-PA-DARPin, 10xHis-1TEL-GGG-DARPin, 10xHis-1TEL-[SR]-DARPin, 10xHis-1TEL-[MR]-DARPin, 10xHis-1TEL-[HLX]-DARPin, and 10xHis-1TEL-[SR]-UBA (without cleavable 10xHis-SUMO tag)

All steps for these constructs were identical to those described above for SUMO-1TEL-PA-DARPin, except the SUMO tag cleavage and tag removal steps were omitted.

### 5.5. Crystallization

The purified proteins’ concentrations were set to 1, 9, and 18 mg/mL and screened against both commercial conditions (PEG Ion, Salt RX, and Index, Hampton Research) and in-house custom conditions (PEG Custom). 1.2 μL of each purified protein was combined with 1.2 μL of each crystallization solution in a sitting drop format (SPT Labtech Mosquito) and equilibrated against a 50 μL reservoir of each crystallization solution via vapor diffusion. Crystals were mounted and passed through 20% glycerol in reservoir solution as a cryoprotectant before snap freezing in liquid nitrogen.

### 5.6. Data collection, reduction, and structure solution

Crystallographic diffraction data were collected remotely at the Stanford Synchrotron Radiation Lightsource (SSRL) beamlines 9-2 and 12-2 at 100 K. Diffraction images were processed autoPROC, which indexed, integrated, scaled, and merged the data. The structures were solved by molecular replacement, using search models for 1TEL and DARPin derived from PDB ID 7N2B (Nawarathnage et al., 2022) to determine initial phases. Final refinement was performed using Phenix Refine, with iterative cycles of model building in Coot and validation to achieve well-refined structures suitable for structural analysis.

Macromolecule production information [Style: IUCr table caption; this style applies table numbering] In the primers, indicate any restriction sites, cleavage sites or introduction of additional residues, *e.g.* His6-tag, as well as modifications, *e.g.* Se-Met instead of Met. [Style: IUCr table headnote]

**Table 6.** Table1 From Each Structure Solution. Values for the outer shell are given in parentheses.

|  | 3TEL-GGG-DARPin (PDB ID: 9E4Q) | 10xHis-1TEL-[MR]-DARPin (PDB ID: 9DP8) | 10xHis-1TEL-[SR]-DARPin '2-fold' crystal form (PDB ID: 9DVG) | 10xHis-1TEL-[SR]-DARPin '3-fold' crystal form (PDB ID: 9DAM) | 1TEL-PA-DARPin (PDB ID: 9DB5) |
| --- | --- | --- | --- | --- | --- |
| Wavelength ($\text{\AA}$) | 0.979460 | 0.979460 | 0.979460 | 0.979460 | 0.979460 |
| **Resolution range**<br>(Å) | 42.75 - 3.58 (3.708<br>- 3.58) | 45.89 - 3.474<br>(3.598 - 3.474) | 43.93 - 3.542<br>(3.668 - 3.542) | 41.65 - 1.78 (1.844<br>- 1.78) | 48.87 - 1.57 (1.626<br>- 1.57) |
| **Space group** | P 2<sub>1</sub> 2<sub>1</sub> 2<sub>1</sub> | P 6<sub>5</sub> | P 6<sub>5</sub> | P 6<sub>5</sub> | P 6<sub>5</sub> |
| **Unit cell** | 46.14 85.5 120.675<br>90 90 90 | 105.981 105.981<br>51.137 90 90 120 | 101.461 101.461<br>48.077 90 90 120 | 83.29 83.29 58.434<br>90 90 120 | 97.73 97.73 45.816<br>90 90 120 |
| **Total reflections** | 75782 (6751) | 37143 (3892) | 73291 (6217) | 464490 (44188) | 452015 (49961) |
| **Unique reflections** | 6006 (586) | 3961 (320) | 3560 (359) | 22160 (1639) | 33052 (3468) |
| **Multiplicity** | 12.6 (11.5) | 9.4 (9.2) | 20.6 (17.3) | 21.0 (20.2) | 13.7 (14.3) |
| **Completeness (%)** | 99.57 (98.82) | 87.47 (75.29) | 99.92 (100.00) | 97.23 (75.01) | 94.10 (99.54) |
| **Mean I/σ(I)** | 8.55 (2.36) | 7.03 (0.50) | 8.12 (1.86) | 14.13 (0.60) | 12.34 (1.38) |
| **Wilson B-factor**<br>(Å<sup>2</sup>) | 80.19 | 147.09 | 135.55 | 30.93 | 17.08 |
| **R-merge** | 0.2825 (1.132) | 0.1269 (3.2) | 0.1917 (2.487) | 0.1516 (5.909) | 0.2162 (4.199) |
| **R-meas** | 0.2945 (1.187) | 0.1343 (3.39) | 0.1966 (2.562) | 0.1553 (6.06) | 0.2246 (4.353) |
| **R-pim** | 0.08171 (0.3483) | 0.04332 (1.111) | 0.04311 (0.6086) | 0.03361 (1.332) | 0.05997 (1.14) |
| **CC 1/2** | 0.993 (0.828) | 0.999 (0.578) | 0.999 (0.407) | 0.999 (0.335) | 0.996 (0.501) |
| **CC\*** | 0.998 (0.952) | 1 (0.856) | 1 (0.761) | 1 (0.708) | 0.999 (0.817) |
| **Reflections used**<br>**in refinement** | 5999 (585) | 3806 (320) | 3559 (359) | 21600 (1639) | 32999 (3467) |
| **Reflections used**<br>**for R-free** | 308 (30) | 189 (21) | 163 (16) | 1054 (76) | 1585 (174) |
| **R-work** | 0.2604 (0.3063) | 0.2651 (0.3778) | 0.2967 (0.4073) | 0.2063 (0.3332) | 0.1671 (0.2556) |
| **R-free** | 0.3267 (0.3946) | 0.2797 (0.3668) | 0.3131 (0.4339) | 0.2263 (0.3629) | 0.1963 (0.2884) |
| **CC (work)** | 0.922 (0.845) | 0.955 (0.604) | 0.907 (0.381) | 0.954 (0.670) | 0.963 (0.797) |
| **CC (free)** | 0.869 (0.504) | 0.906 (0.710) | 0.780 (0.384) | 0.945 (0.622) | 0.962 (0.786) |
| **Number of non-**<br>**hydrogen atoms** | 2638 | 1657 | 1748 | 1918 | 2013 |
| **Macromolecules** | 2611 | 1655 | 1748 | 1786 | 1792 |
| **Ligands** | 15 | 0 | 0 | 12 | 100 |
| **Solvent** | 12 | 2 | 0 | 123 | 160 |
| **Protein residues** | 373 | 233 | 228 | 238 | 234 |
| **RMS (bonds)** | 0.003 | 0.006 | 0.004 | 0.012 | 0.011 |
| **RMS (angles)** | 0.5 | 0.78 | 0.67 | 1.12 | 1.05 |
| **Ramachandran**<br>**favored (%)** | 96.46 | 93.07 | 96.90 | 98.72 | 99.14 |
| **Ramachandran**<br>**allowed (%)** | 3.54 | 6.93 | 3.10 | 0.43 | 0.86 |
| **Ramachandran**<br>**Outliers (%)** | 0.00 | 0.00 | 0.00 | 0.85 | 0.00 |
| **Rotamer outliers**<br>**(%)** | 0.00 | 0.00 | 0.57 | 0.59 | 0.00 |
| **Clashscore** | 8.41 | 12.16 | 8.19 | 3.79 | 3.87 |
| **Average B-factor**<br>**(Å<sup>2</sup>)** | 68.78 | 211.57 | 156.62 | 36.09 | 22.36 |
| **Macromolecule** | 68.71 | 211.79 | 156.62 | 35.55 | 21.07 |
| **Ligands** | 100.52 | 0 | 0 | 55.83 | 33.39 |
| **Solvent** | 45.11 | 30.00 | 0 | 42.44 | 32.57 |
| **Number of TLS**<br>**groups** | 3 | NULL | NULL | NULL | 6 |

## Acknowledgements

Use of the Stanford Synchrotron Radiation Lightsource, SLAC National Accelerator Laboratory, is supported by the U.S. Department of Energy, Office of Science, Office of Basic Energy Sciences under Contract No. DE-AC02-76SF00515. The SSRL Structural Molecular Biology Program is supported by the DOE Office of Biological and Environmental Research, and by the National Institutes of Health, National Institute of General Medical Sciences (including P41GM103393). The contents of this publication are solely the responsibility of the authors and do not necessarily represent the official views of NIGMS or NIH. Research reported in this publication was supported by the National Institute of General Medical Sciences of the National Institutes of Health under award numbers R15GM146209 and R35GM155011. The content is solely the responsibility of the authors and does not necessarily represent the official views of the National Institutes of Health.

---

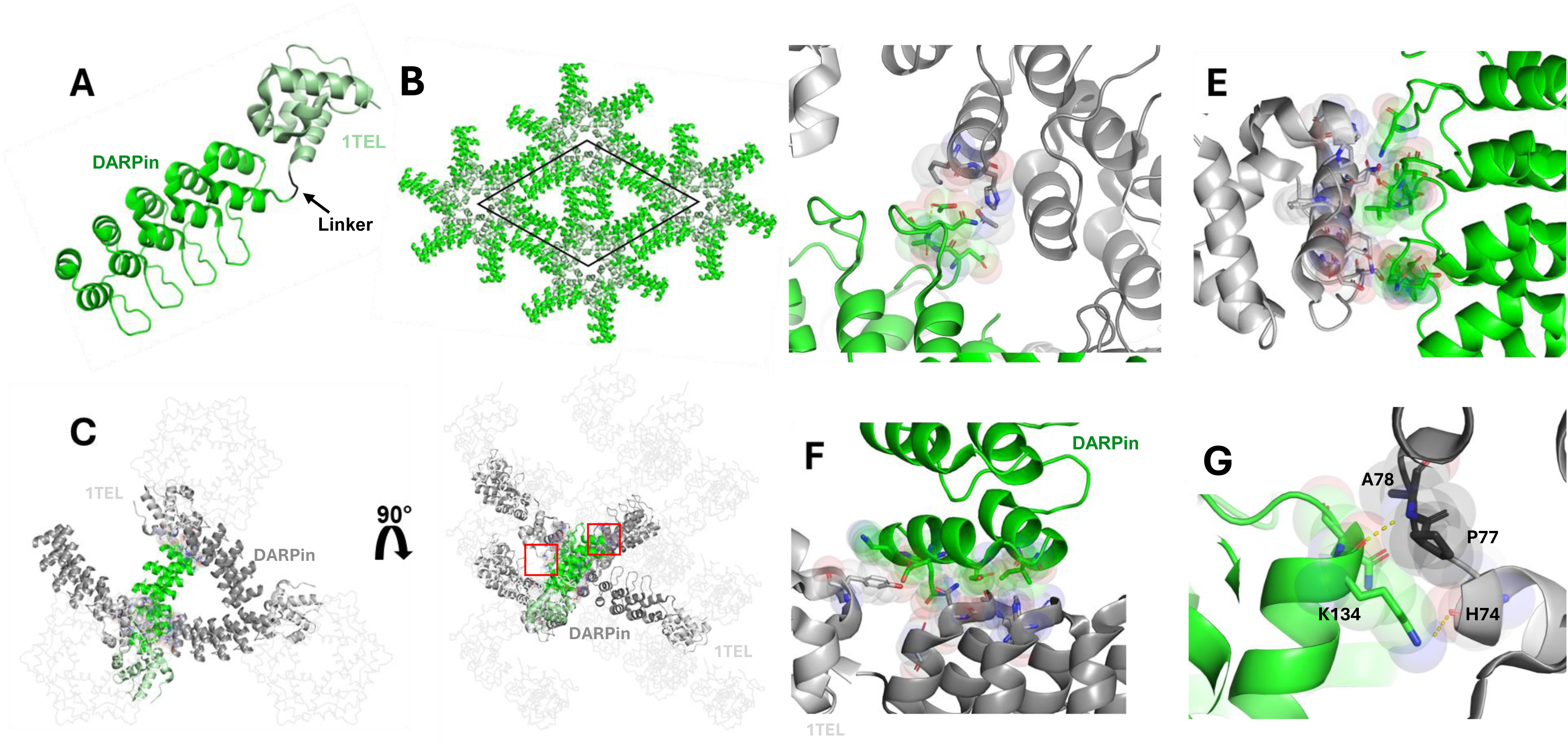

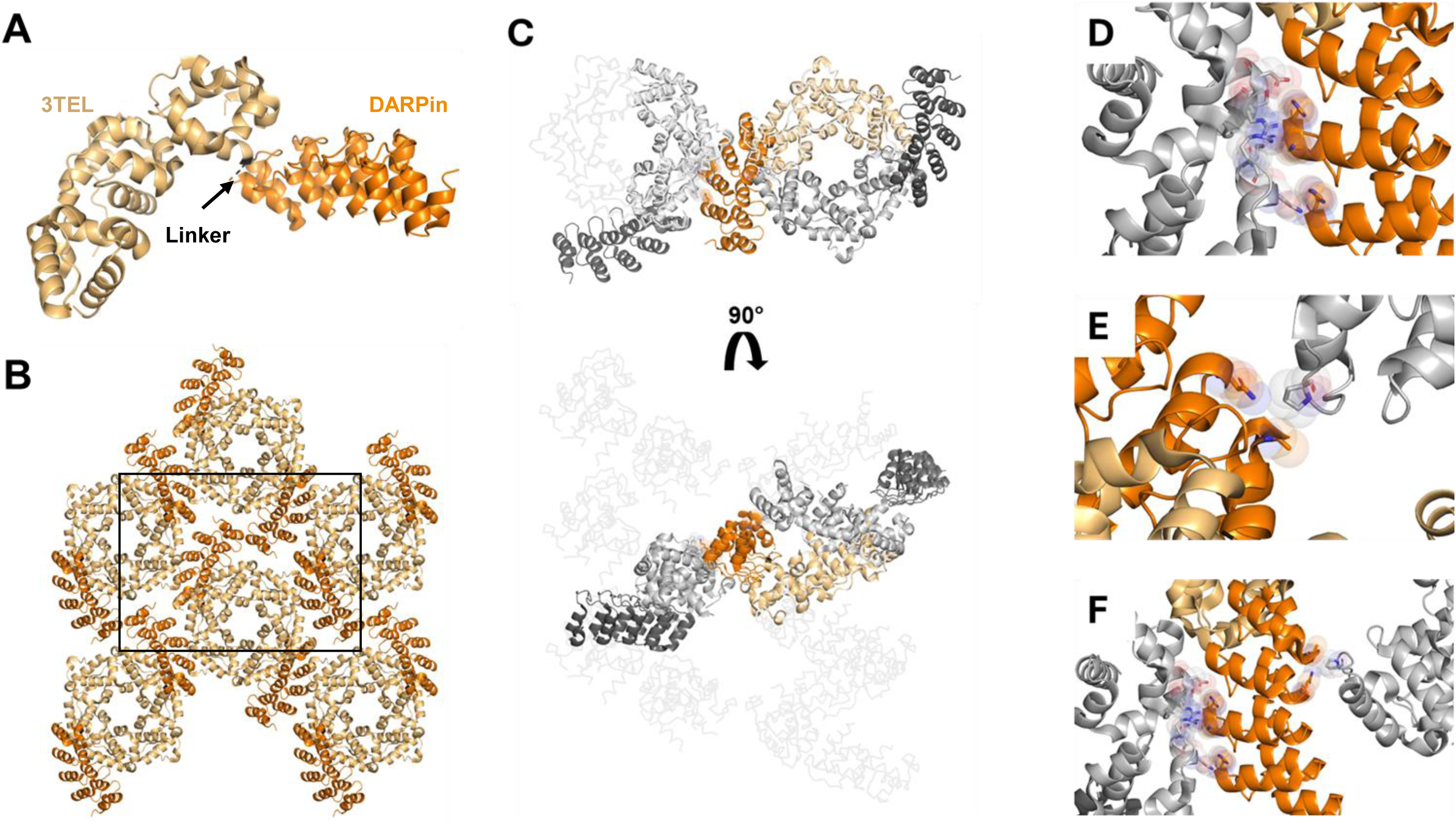

## References

1. Wei, H.; McCammon, J. A. Structure and dynamics in drug discovery. npj Drug Discovery 2024, 1, 1 DOI:10.1038/s44386-024-00001-2.

2. Ferreira, L. G.; Dos Santos, R. N.; Oliva, G.; Andricopulo, A. D. Molecular docking and structure-based drug design strategies. Molecules 2015, 20, 13384–13421 DOI:10.3390/molecules200713384.

3. Lionta, E.; Spyrou, G.; Vassilatis, D. K.; Cournia, Z. Structure-based virtual screening for drug discovery: principles, applications and recent advances. Curr. Top. Med. Chem. 2014, 14, 1923–1938 DOI:10.2174/1568026614666140929124445.

4. Maveyraud, L.; Mourey, L. Protein X-ray Crystallography and Drug Discovery. Molecules 2020, 25 DOI:10.3390/molecules25051030.

5. Dale, G. E.; Oefner, C.; D’Arcy, A. The protein as a variable in protein crystallization. J. Struct. Biol. 2003, 142, 88–97 DOI:10.1016/S1047-8477(03)00041-8.

6. Terwilliger, T. C.; Stuart, D.; Yokoyama, S. 2. Lessons from Structural Genomics*. Annual Review of Biophysics,, 38, 371–383 DOI:10.1146/annurev.biophys.050708.133740.

7. Cooper, D.,R.;, Porebski, Przemyslaw, J.; Chruszcz, M.; and Minor, W. X-ray crystallography: assessment and validation of protein–small molecule complexes for drug discovery. Expert Opinion on Drug Discovery 2011, 6, 771–782 DOI:10.1517/17460441.2011.585154.

8. Derewenda, Z. S. Application of protein engineering to enhance crystallizability and improve crystal properties. Acta Crystallogr. D Biol. Crystallogr. 2010, 66, 604–615 DOI:10.1107/S090744491000644X.

9. Nauli, S.; Farr, S.; Lee, Y.; Kim, H.; Faham, S.; Bowie, J. U. Polymer-driven crystallization. Protein Science 2007, 16, 2542–2551 DOI:10.1110/ps.073074207.

10. Gajjar, P. L.; Pedroza Romo, M. J.; Litchfield, C. M.; Callahan, M.; Redd, N.; Nawarathnage, S.; Soleimani, S.; Averett, J.; Wilson, E.; Lewis, A.; Stewart, C.; Tseng, Y.; Doukov, T.; Lebedev, A.; Moody, J. D. Increasing the bulk of the 1TEL--target linker and retaining the 10$\times$His tag in a 1TEL--CMG2-vWa construct improves crystal order and diffraction limits. Acta Crystallographica Section D 2023, 79, 925–943 DOI:10.1107/S2059798323007246.

11. Nawarathnage, S.; Tseng, Y. J.; Soleimani, S.; Smith, T.; Pedroza Romo, M. J.; Abiodun, W. O.; Egbert, C. M.; Madhusanka, D.; Bunn, D.; Woods, B.; Tsubaki, E.; Stewart, C.; Brown, S.; Doukov, T.; Andersen, J. L.; Moody, J. D. Fusion crystallization reveals the behavior of both the 1TEL crystallization chaperone and the TNK1 UBA domain. Structure 2023, 31, 1589–1603.e6 DOI:10.1016/j.str.2023.09.001.

12. Nawarathnage, S.; Soleimani, S.; Mathis, M. H.; Bezzant, B. D.; Ramírez, D. T.; Gajjar, P.; Bunn, D. R.; Stewart, C.; Smith, T.; Pedroza Romo, M. J.; Brown, S.; Doukov, T.; Moody, J. D. Crystals of TELSAM–target protein fusions that exhibit minimal crystal contacts and lack direct inter-TELSAM contacts. Open Biology 2022, 12, 210271 DOI:10.1098/rsob.210271.

13. McFedries, A.; Schwaid, A.; Saghatelian, A. Methods for the Elucidation of Protein-Small Molecule Interactions. Chem. Biol. 2013, 20, 667–673 DOI:10.1016/j.chembiol.2013.04.008.

14. Seeger, M. A.; Zbinden, R.; Flütsch, A.; Gutte, P. G. M.; Engeler, S.; Roschitzki-Voser, H.; Grütter, M. G. Design, construction, and characterization of a second-generation DARPin library with reduced hydrophobicity. Protein Science 2013, 22, 1239–1257 DOI:10.1002/pro.2312.

15. Kottur, J.; Rechkoblit, O.; Quintana-Feliciano, R.; Sciaky, D.; Aggarwal, A. K. High-resolution structures of the SARS-CoV-2 N7-methyltransferase inform therapeutic development. Nature Structural & Molecular Biology 2022, 29, 850–853 DOI:10.1038/s41594-022-00828-1.

16. Poulos, S.; Agah, S.; Jallah, N.; Faham, S. Symmetry based assembly of a 2 dimensional protein lattice. PLOS ONE 2017, 12, e0174485.

17. Kleffner, R.; Flatten, J.; Leaver-Fay, A.; Baker, D.; Siegel, J. B.; Khatib, F.; Cooper, S. Foldit Standalone: a video game-derived protein structure manipulation interface using Rosetta. Bioinformatics 2017, 33, 2765–2767 DOI:10.1093/bioinformatics/btx283.

18. Walls, W. G.; Moody, J. D.; McDaniel, E. C.; Villanueva, M.; Shepard, E. M.; Broderick, W. E.; Broderick, J. B. The B12-independent glycerol dehydratase activating enzyme from Clostridium butyricum cleaves SAM to produce 5′-deoxyadenosine and not 5′-deoxy-5′-(methylthio)adenosine. J. Inorg. Biochem. 2022, 227, 111662 DOI:10.1016/j.jinorgbio.2021.111662.

19. Gibson, D. G.; Young, L.; Chuang, R.; Venter, J. C.; Hutchison, C. A.; Smith, H. O. Enzymatic assembly of DNA molecules up to several hundred kilobases. Nature Methods 2009, 6, 343–345 DOI:10.1038/nmeth.1318.

20. Lau, Y. K.; Baytshtok, V.; Howard, T. A.; Fiala, B. M.; Johnson, J. M.; Carter, L. P.; Baker, D.; Lima, C. D.; Bahl, C. D. Discovery and engineering of enhanced SUMO protease enzymes. J. Biol. Chem. 2018, 293, 13224–13233 DOI:10.1074/jbc.RA118.004146.

